# More than the sum of its parts: Merging network psychometrics and network neuroscience with application in autism

**DOI:** 10.1101/2020.11.17.386276

**Authors:** Joe Bathelt, Hilde M. Geurts, Denny Borsboom

**Affiliations:** Royal Holloway, University of London, Department of Psychology, Egham, Surrey, TW20 0EX, United Kingdom; University of Amsterdam, Department of Psychology, Amsterdam, 1018 WV, The Netherlands

## Abstract

Network approaches that investigate the interaction between symptoms and behaviours have opened new ways of understanding psychological phenomena in health and disorder in recent years. In parallel, network approaches that characterise the interaction between brain regions have become the dominant approach to understanding brain function in neuroimaging research. Combining these parallel approaches would enable new insights into the interaction between behaviours and their brain-level correlates. In this paper, we introduce a methodology for combining network psychometrics and network neuroscience. This approach utilises the information from the psychometric network to obtain neural correlates that are associated with each node in the psychometric network (network-based regression). Moreover, we combine the behavioural variables and their neural correlates in a joint network to characterise their interactions. We illustrate the approach by highlighting the interaction between the triad of autistic traits and their resting-state functional connectivity associations. To this end, we utilise data from 172 male autistic participants (10-21 years) from the autism brain data exchange (ABIDE, ABIDE-II) that completed resting-state fMRI and were assessed using the autism diagnostic interview (ADI-R). Our results indicate that the network-based regression approach can uncover both unique and shared neural correlates of behavioural measures. In addition, because the shared variance between behavioural measures is controlled for in the approach, the methodology enables us to isolate mechanisms at the brain-level that are unique to particular behavioural variables. For instance, our example analysis indicates that the overlap between communication and social difficulties is not reflected in the overlap between their functional brain correlates.

## 1 Introduction

The traditional view of psychiatric conditions conceptualizes mental disorders as latent constructs that are expressed in a set of manifest symptoms. For instance, autism spectrum disorders (ASD) are typically described as a disorder rooted in a (as yet unknown) dysfunction in the biology of the human system, which manifests its effects in the domains of social interaction, social communication, and restricted, repetitive patterns of behaviour, interests, or activities [DSM-5]. In this view, the manifested difficulties – typically interpreted as symptoms – are viewed as indicators of a latent condition; in accordance, the information present in these symptoms is commonly aggregated into a single score (e.g. by counting the number of symptoms or averaging subscales of questionnaires). This score is interpreted as a measure of the severity of the mental disorder, and usually functions as a dependent variable in designs geared to uncover the genetic background or neural correlates of the disorder.

An alternative view, which has emerged in recent years, interprets the relation between symptoms and disorders differently, and emphasises the dynamic interaction between symptoms^1–3^. In this alternative network approach to mental disorders, causal interaction between symptoms themselves is brought to the forefront and disorders are seen as clustered states of symptoms that mutually reinforce each other. For instance, in the case of autism, restricted interests may limit the time spent in social interactions, which may lead to reduced social communication skills, and the limited success in social communication may in turn reinforce restricted interests. From this perspective, an important part of the basis of mental disorders like autism would be expected to lay in the disposition to develop individual symptoms and in the processes that govern interactions between them. As a result, the network approach does not focus on the level of aggregate scores or functions thereof, but on the patterns of association that arise between symptom variables. A host of statistical techniques has been developed to estimate and analyse such symptom networks^4^. This way of working has become a popular methodology approach to the study of symptomatology and is beginning to find its way in clinical practice^5^.

By focusing attention on the interaction between symptoms, network approaches open up new ways to study how mental disorders may be related to the brain. In particular, from the point of view of network theory, neuroscientific approaches should focus on (a) mechanisms or dysfunctions that lead symptoms to arise (e.g., mechanisms that promote repetitive behaviours), and (b) processes that couple one symptom to another (e.g., mechanisms that link repetitive behaviours to, say, problems in social interaction). To the extent that biological processes are specific to individual symptoms or symptom-symptom interactions, aggregation of scores is methodologically inadvisable because the shared variance between symptoms is then likely to correspond to a biological amalgam of processes and mechanisms that will be hard to tease apart.

Methodologies developed in the network approach may be useful in disentangling neural correlates of symptoms and interactions between them. However, a clear link between the network psychometrics approach and brain imaging research has so far been missing. Establishing this connection has the potential to open a new line of research that investigates the relations between brain-level and symptom-level networks and, thereby, holds the potential to uncover mechanisms of compensation or vulnerability. However, the current approaches in psychiatric neuroimaging are not well suited to this task. The dominant approaches are to either contrast groups of cases with groups of controls, or to identify correlates of symptoms or putative cognitive endophenotypes. Both approaches have limitations as has been discussed at length elsewhere^6,7^. Most importantly for network psychometrics, these approaches implicitly assume that psychiatric conditions are latent constructs that determine and, therefore, are indicated by symptoms or cognitive endophenotypes. In this manuscript, we introduce an alternative approach based on network psychometrics that uses information from the symptom network to create network-based regressors. Subsequently, we explore the relationship between the symptom-level network and its neural correlates. For this purpose, we focus on the triad of autistic traits. ASD provides an ideal test case for the application of the combined network psychometric and network neuroscience approach, because of the large literature regarding the association of behavioural measures and their genetic and neural correlates (see Supplementary Introduction for an overview of this literature).

## 2 Methods

### 2.1 Network-based regression method

The following section describes new methods for constructing networks based on brain correlates of behavioural or cognitive measures.

#### 2.1.1 Behaviour-network-based regressors

In neuroimaging, out-of-scanner task performance scores are often used to identify the neural correlates of behaviours, particularly for analyses of resting-state data (fMRI, M/EEG) or structural data (diffusion weighted imaging, morphological analyses). However, the use of raw task scores has been criticised, because the task scores reflect many extraneous influences. One way to create better regressors is to calculate the shared variance of several measures that are thought to tap the same psychological construct^8^. This approach of creating latent variables views the psychological or psychiatric constructs as latent entities indicated by the scores. In contrast, in network psychometrics, one may be more interested in the unique variance of each measure to create the nodes that make up the network. For neuroimaging, this can be achieved by simply regressing the effect of other variables from each variable and retaining the residual. For instance, in a network of three variables A, B, and C, the unique variance in A is given by the residual term, epsilon, in the regression equation: *y_A_* = *β_B_X_B_* + *β_C_X_C_* + *X_Intercept_* + *ε*. The residual terms can be used as regressors to obtain the neural correlates of the unique variance of each behavioural measure.

#### 2.1.2 Associations between behaviour scales and functional brain connectivity

To identify the associations between functional connectivity and behavioural ratings, we used the connectome predictive modelling (CPM) approach described by Shen et al.^9^ (see Figure 1 for an overview). In short, a set of edges is identified that correlate with the behaviour ratings below a certain *p*-value threshold. Then, the edge weights are summed into a brain score and entered into a regression model to estimate the association between the brain score and behaviour ratings. For the current analysis, we identified positively and negatively associated edges separately and entered the summed edge weight for positively and negatively associated edges into a common multiple regression model: *y_symptom_* = *β_Brain^+^_X_Brain^+^_ + β_Brain^-^_X_Brain^-^_ + X_Intercept_* + *ε*. We were faced with multiple methodological choices that were difficult to determine *α priori*, namely the optimal number of ROIs in the parcellation, the edge definition, *p*-value threshold, and global signal regression strategy. Therefore, we employed shuffle split cross-validation to find the combination of parameters that led to the best prediction of behaviour ratings in unseen data. For this parameter tuning, we randomly split the data into an 80% training and 20% test set (see last step in Figure 1). We identified the associated edges and fitted the regression model in the training set. Then, we compared the quality of prediction using the model using the held-out test data by calculating the correlation between predicted and observed values^9^. The parameter combination that produced the best prediction in unseen data used 300 ROIs, defined the edge weight through Pearson correlation without global signal regression applied, and set the association p-value threshold to p<0.001 (see Supplementary Methods for detailed parameter tuning results). For the evaluation of the final model, we only retained edges that were included across 7 out of 10 splits of the data. We compared the performance of this model against 5,000 random permutations of the behavioural data.

**Figure 1.**
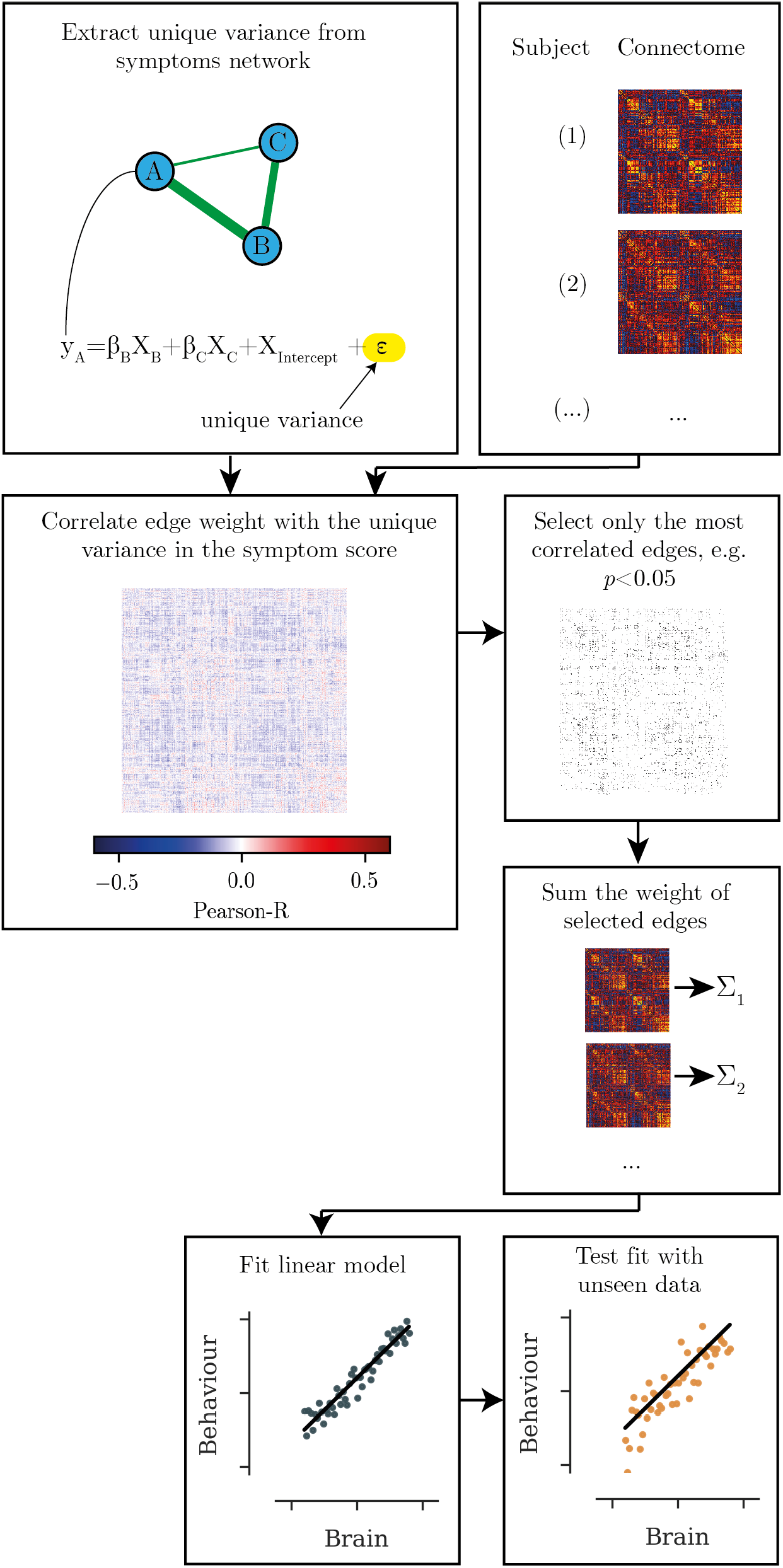
Overview of the analysis steps. First, the unique variance in each behavioural scale is calculated. The edges that shows the strongest correlation with the unique variance in the behavioural score are extracted. The summed edge weight of the most highly associated edges is used to build a regression model, which is then tested in unseen data.

#### 2.1.3 Construction of the behaviour and brain score network

To assess the relations between behavioural scores and their neural correlates, we estimated the network structure of a network based on the behavioural scores and the total brain score for each behavioural measure. To construct the total brain score, the regression weights were applied to the positive and negative brain scores. To determine the edges of the network, we identified an unregularized Gaussian graphical model by minimizing the extended Bayesian Information Criterion (BIC) using the glasso algorithm and stepwise model selection. These analyses were performed in R (version 3.5.0) using the *bootnet* v1.4.3 and *qgrαph* v1.6.5 packages^4^.

Subsequently, we identified the likely causal direction of the network edges^10^ in the network of behavioural scores, rsfMRI correlates, and a combined network with behavioural and rsfMRI correlate nodes. For this purpose, we computed a Bayesian network, described in a directed acyclic graph (DAG), using structure learning algorithms implemented in the bnlearn v4.5 package^11^. The structure learned by these algorithms can contain directed and undirected edges. We compared the structure learned through different algorithms, namely the *Grow-Shrink^12^, Incremental Association^13^, Fast Incremental Association^14^, Interleaved Incremental Association*^13^, and *Max-Min Parents and Children*^15^ algorithms. We also compared the structure learned when using different independence test statistics, namely *Pearson correlation, Fisher’s Z, Monte-Carlo permutation*, and *mutual information*.

### 2.2 General Data Preparation

The following section describes the data and pre-processing steps that were used to illustrate the network-based regression approach.

#### 2.2.1 Participants

The analysis was based on data taken from the first and second Autism Brain Imaging Data Exchange database (ABIDE^16^ and ABIDE-II^17^). Both databases collated resting-state fMRI and phenotypic data from autistic participants from 19 international sites. There was no prior coordination between sites, which means that the sites differed in their fMRI acquisition protocols and diagnostic procedures. Because of the ensuing variability, we applied selection criteria to arrive at a more homogeneous sample. Namely, we selected only male participants, because women were not well represented. Further, we selected only participants over 10 years of age, because of different scoring criteria on the Autism Diagnostic Interview - Revised (ADI-R)^18^ in younger children (see below for a description of the ADI-R). We further excluded participants older than 21 years, because the relatively few participants older than 21 years were spread over a large age range. In addition, we only selected participants with complete, research-reliable ADI-R assessments and with complete structural and functional MRI that was rated as useable by expert human assessors [Note: The quality ratings are distributed with the phenotypic data]. For further quality control, we excluded 13 participants because their fMRI data fell outside the recommended range on established quality metrics (framewise displacement > 0.5mm^19^, DVARS > 5%^20^, see Supplementary Methods for further details regarding quality control). The final sample consisted of 172 autistic participants (ABIDE: n=127, ABIDE-II: n=45, see Table for sample characteristics). Please note that were not aiming to obtain a representative sample. The purpose of the analysis was to demonstrate the potential use of a network-based regression method. The selection criteria were intended to create a more homogeneous sample with good quality imaging data.

#### 2.2.2 Assessment of autism characteristics

The Autism Diagnostic Interview-Revised (ADI-R) is a standardised diagnostic interview for primary caregivers^18^. It focuses on a description of a child’s behaviour when they were 4-5 years old and their current behaviour. An autism diagnosis is made with an ADI-R algorithm that consists of 37 extracted items. In addition to a total score, sub-scores for autistic traits (social interaction, communication, RRBI) can be obtained. The ADI-R shows a higher inter-rater agreement (0.94-0.96)^21^ and high convergence with clinical team assessments and another commonly used assessment protocol (75% agreement)^22^, i.e. the autism observation schedule (ADOS). The ADI-R protocol is adjusted depending on the chronological and mental age of the participant. Fewer items are included for children younger than 10 years or with an intellectual functioning outside of the typical range. Because the current analysis aimed to compare associations of scores, we restricted our analysis to children above 10 years with a full-scale IQ in the typical range to ensure that differences in ADI-R scores reflected the relative ranking of the severity of difficulties in each domain.

#### 2.2.3 fMRI processing

The current analysis was based on data processed using a standardised state-of-the-art pipeline for resting-state fMRI processing for functional connectomics^23^ (see Supplementary Methods for full details of fMRI processing). We used cross-validation to make data-informed decision regarding the granularity of the parcellation, definition of the connectome edges, and sparsity threshold for the brain-behaviour associations. The full details of the parameter tuning can be found in the Supplementary Methods.

#### 2.2.4 Interpretation of brain-behaviour associations

To aid the objective interpretation of the brain areas found to be associated with behaviour ratings using the connectome predictive modelling method, we utilised the NeuroSynth meta-analytic engine^24^. NeuroSynth collects reported activations from thousands of published articles and associates them with terms used in the description of the results. This leads to meta-analytic maps that associate brain activations with cognitive descriptors. NeuroSynth also includes a decoding feature that finds cognitive terms with brain maps that most closely resemble a map uploaded to NeuroSynth. We made use of this feature here. Namely, we converted the matrix of edges that were consistently associated with the behaviour rating scale to a brain map by summing all edges associated with a node and assigning the resulting value to that ROI in the parcellation. All other values were set to 0. Next, we min-max-scaled the values to get all images into the same value range. We then used the decoding feature of NeuroSynth to obtain a set of cognitive terms that showed the best correspondence with the brain-behaviour association maps. We evaluated the 25 most highly associated terms for each map and disregarded any anatomical or descriptive terms, e.g. “visual cortex” or “default mode network”. We removed highly similar terms and retained the term that was most strongly associated, e.g. ‘autobiographical’ vs ‘autobiographical memory’. The included terms were: ‘affective’, ‘autobiographical’, ‘competition’, ‘facial expression’, ‘emotion’, ‘error’, ‘executive’, ‘moral’, ‘painful’, ‘theory [of] mind’, ‘working memory’. Subsequently, we calculated the correlation between the meta-analytic map for each of the cognitive terms with the normalised brain-behaviour association maps to assess the strength of the correspondence.

## 3 Results

The connectome-based predictive modelling methods indicated a combination of edge weights that could predict behaviour scores in held-out data (Social: mean=0.14, SE=0.054; Communication: mean=0.05, SE=0.043, RRBI: mean=0.23, SE=0.024, mean correlation between observed and predicted scores across 10 random 80/20 splits) and were significantly better in predicting behaviour scores compared to scrambled data (Social: *p*=0.021; Communication: *p*=0.014, RRBI: *p*=0.011; *p*-value based on 5,000 permutations).

For social difficulties, connections between the limbic (Limbic) and frontoparietal control network (Cont), and connections between the visual and salience/ventral attention network (SalVentAttn) were most strongly related to higher difficulty ratings (see Figure 2 A & C). In contrast, connections of Cont with SalVentAttn, SomMot, and connection between Vis and the default mode network (Default) were related to lower social difficulty ratings (see Figure 2 B & C). Regarding the interpretation of the involved regions, the pattern of edges that was negatively associated with social difficulty scores matched best with the meta-analytic maps for the terms ‘executive’ and ‘working memory’ (see Figure 2 E). The patterns of edges that were related to higher social difficulty scores were most similar to the meta-analytic maps for the terms ‘facial expression’, ‘executive’, and ‘working memory’ (see Figure 2 E).

**Figure 2.**
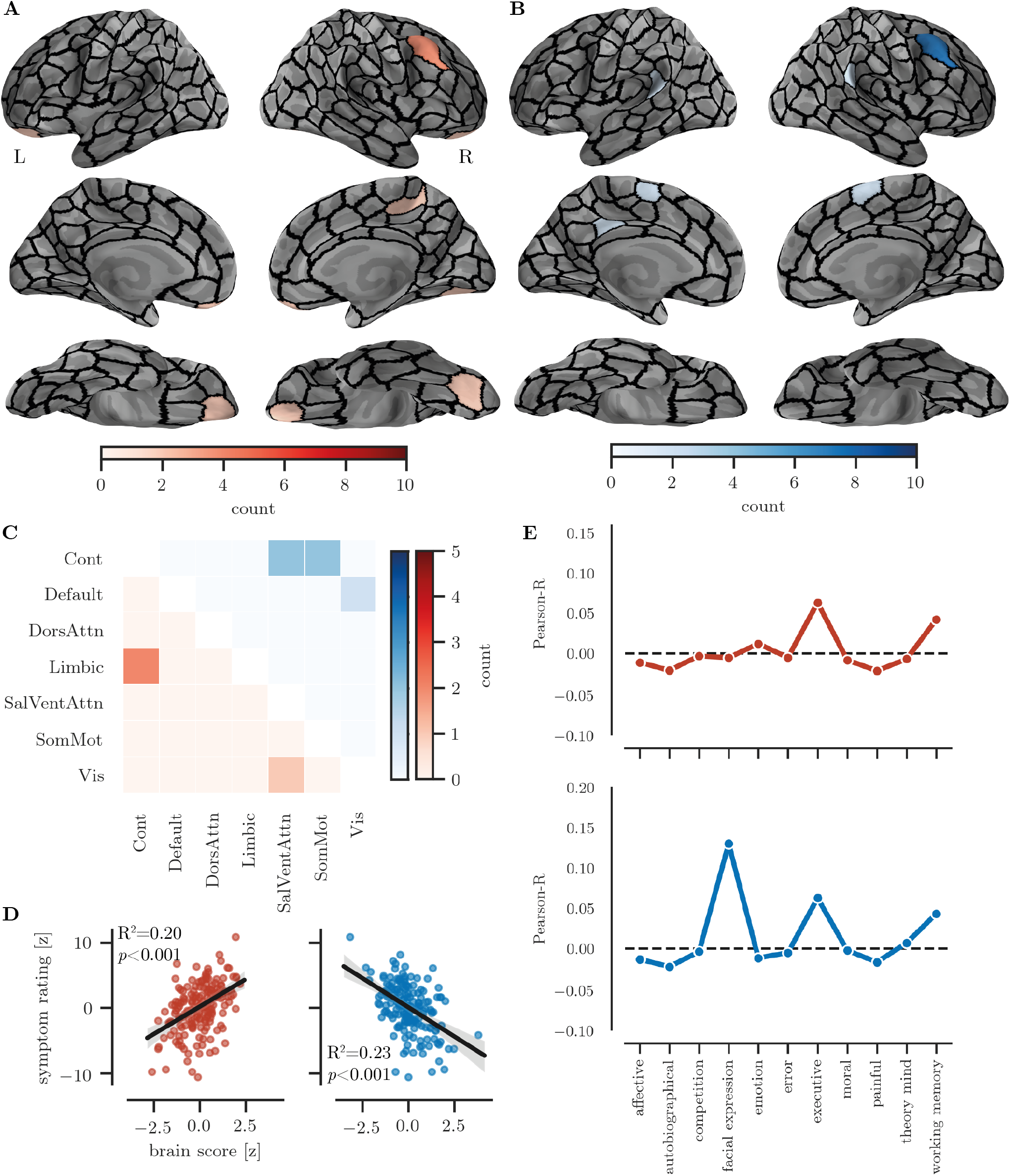
Illustration of the novel brain-behavior-network approach in autistic individuals. i.e, association between functional connectivity and ADI-R Social Scores. **A** Nodes with connections that were consistently positively associated with ADI-R Social scores. **B** Nodes with connections that were consistently negatively associated with ADI-R Social scores. **C** Connections summed by large-scale canonical network. The lower triangle shows positive associations and upper triangle shows negative associations. **D** Association between the summed brain score and the ADI-R Social score. **E** Strength of the association between the connection map and NeuroSynth meta-analytic maps for each cognitive term.

For communication scores, connections between Cont and SomMot, and Cont and Limbic, as well as connections between Default and Vis were associated with higher communication difficulty scores (see Figure 3 A & B). In contrast, connections between regions of the Cont and Vis, and Default and Vis were related to lower communication difficulty ratings (see Figure 3 B). Regarding the interpretation of the involved regions, the pattern of edges that were related to higher communication difficulty were most associated with the meta-analytic maps for the terms ‘error’ and ‘executive’ (Figure 3 E). There were only weak associations for the pattern of edges that were related to lower communication difficulties. The strongest associations were for the terms ‘affective’ and ‘working memory’ (see Figure 3 E)

**Figure 3.**
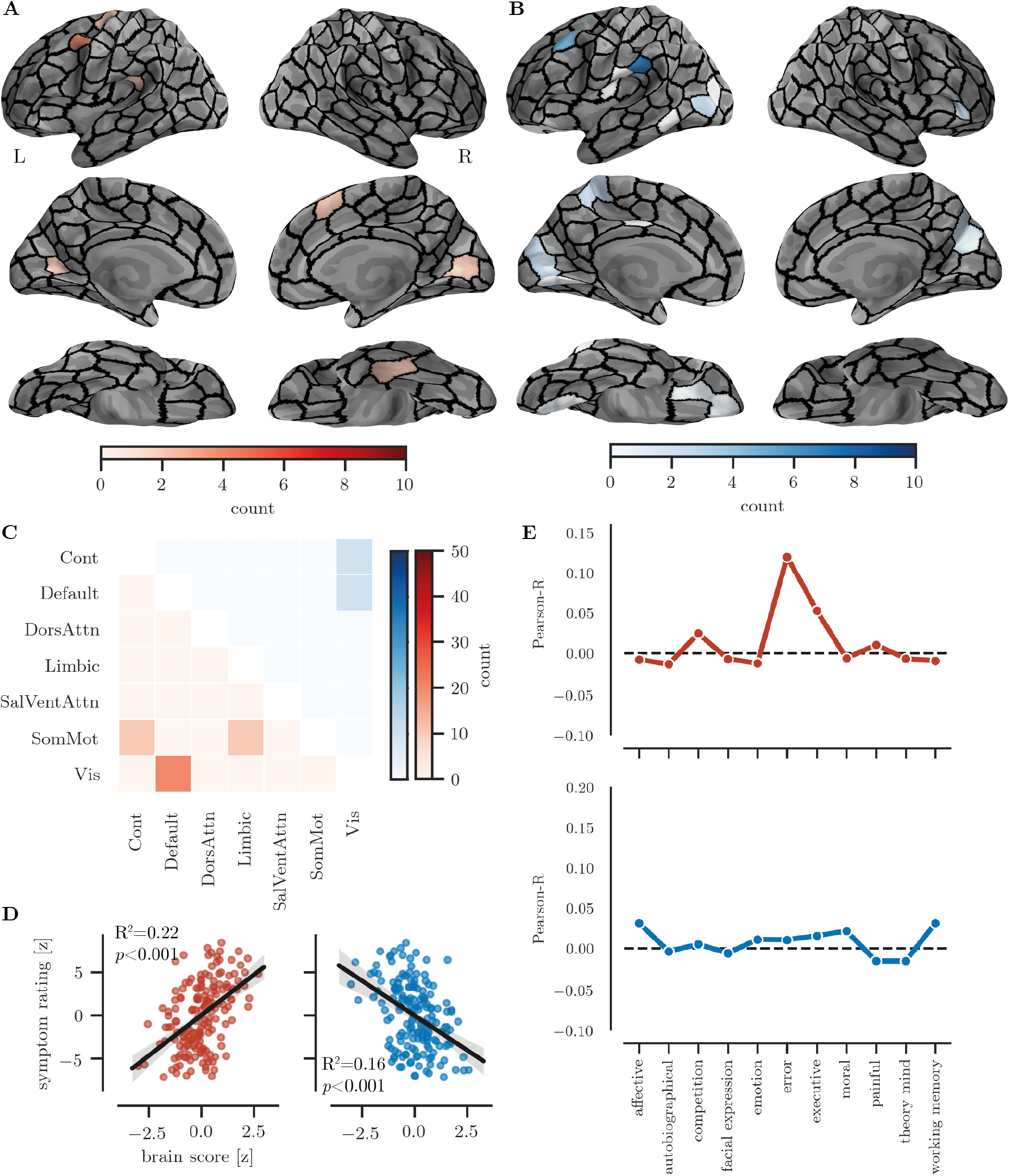
Illustration of the novel brain-behavior-network approach in autistic individuals, i.e, association between functional connectivity and ADI-R Communication Scores. **A** Nodes with connections that were consistently positively associated with ADI-R Verbal scores. **B** Nodes with connections that were consistently negatively associated with ADI-R Communication scores. **C** Connections summed by large-scale canonical network. The lower triangle shows positive associations and upper triangle shows negative associations. **D** Association between the summed brain score and the ADI-R Communication score. **E** Strength of the association between the connection map and NeuroSynth meta-analytic maps for each cognitive term.

For RRBI scores, connections between Cont and Limbic, and Vis and SalVentAttn were related to higher RRBI difficulty ratings (see Figure 4 A & C). Connections of Cont with SalVentAttn and SomMot and connections of Default and Vis were related to lower RRBI difficulty ratings (see Figure 4 B & C). Regarding the interpretation of the involved edges, the pattern of edges that were associated with higher RRBI difficulty ratings was most similar to the meta-analytic maps for the term ‘painful’. The terms most associated with the pattern of edges that were related to lower difficulty ratings were most strongly related to the meta-analytic maps for the terms ‘autobiographical’, and ‘theory [of] mind’.

**Figure 4.**
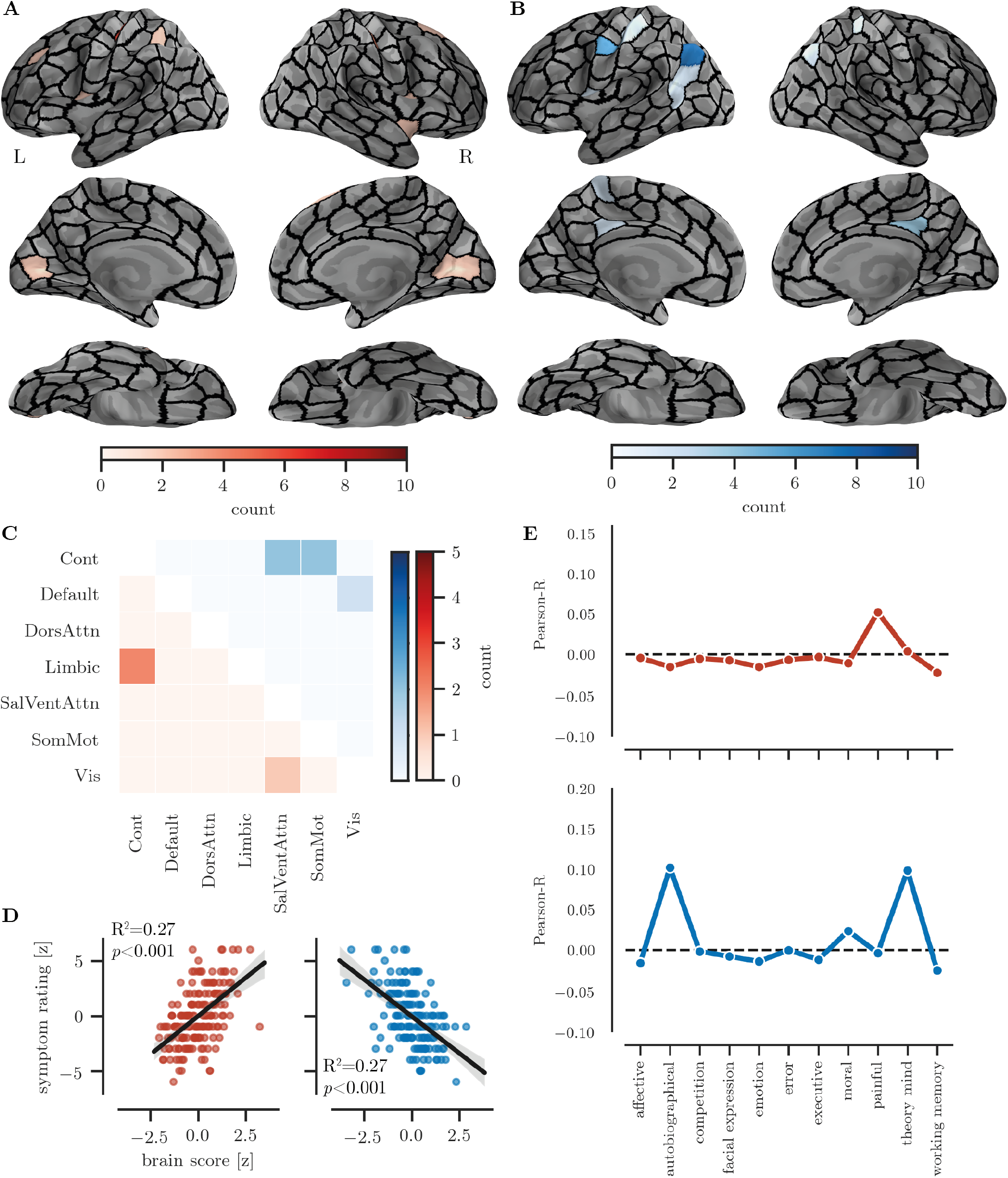
Illustration of the novel brain-behavior-network approach in autistic individuals, i.e. association between functional connectivity and ADI-R RRBI Scores). **A** Nodes with connections that were consistently positively associated with ADI-R RRBI scores. **B** Nodes with connections that were consistently negatively associated with ADI-R Social scores. **C** Connections summed by large-scale canonical network. The lower triangle shows positive associations and upper triangle shows negative associations. **D** Association between the summed brain score and the ADI-R RRBI score. **E** Strength of the association between the connection map and NeuroSynth meta-analytic maps for each cognitive term.

### 3.1 Combined behaviour and brain network

We compared the network structure of a network based on behavioural measures and a network based on the neural correlates of the behavioural measures (see Table 2 for correlations between all measures). As expected from previous studies, the behavioural network showed a strong association between Communication and Social Scores, and a weaker association between Social and RRBI scores (see Figure 5 A). In contrast, the network of rsfMRI correlates showed a strong association between the functional brain correlates of Social and RRBI scores, but no association between the correlates of Social and Communication scores (see Figure 5 B).

**Figure 5.**
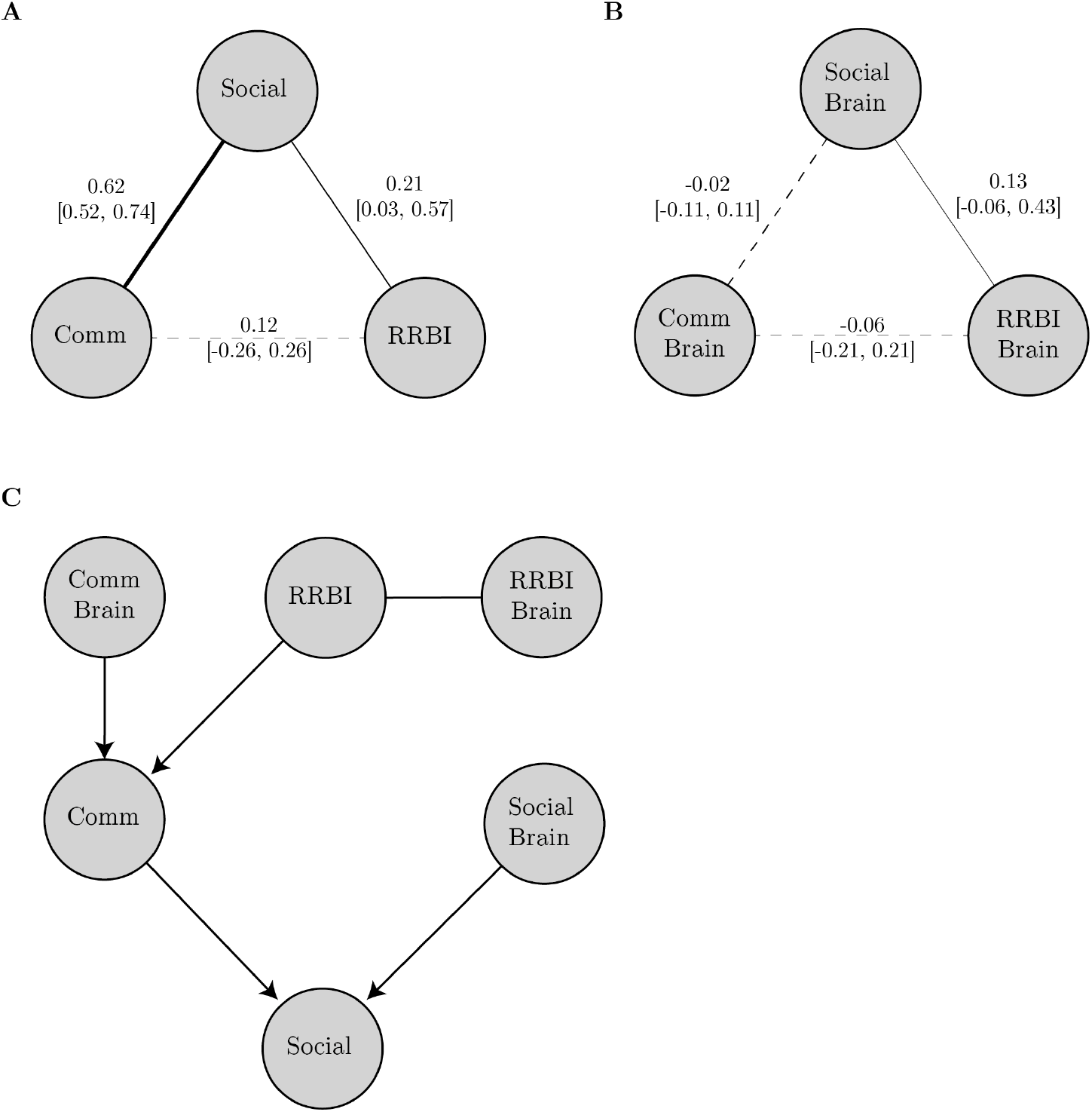
Overview of the network analysis **A & B**: Bootstrapped undirected network structure (A: behaviour, B: rsfMRI correlates). Solid lines indicate significant edges, dashed lines indicate non-significant ones. The thickness of the lines indicates the strength of the association. All edges are positive. The numbers indicate the bootstrapped mean with the 5% and 95%ile in brackets. **C**: Directed acyclic graph determined through Bayesian network analysis with constrained-based learning algorithms for the combined brain-behaviour network. Arrows indicated directed interactions. Lines indicate undirected interactions.

Next, we conducted a causal inference analysis using Bayesian networks. For the network of behaviour scores, there were undirected interactions between all behaviour domains, i.e. Social, Communication, and RRBI. This structure was indicated across all structure learning algorithms and statistics for the independence tests. For the network of brain correlates, a structure with an undirected interaction between rsfMRI correlates of Social Scores and RRBI scores was indicated. There were no connections with the rsfMRI correlates of the Communication score. This structure was consistent across structure learning algorithms and independence test statistics. When combining the behaviour and brain scores in one causal inference network, a more complex network structure emerged (see Figure 5 C). In this network, RRBI behavioural scores predicted Communication behaviour scores, which in turn predicted Social behaviour scores. Social and Communication behaviour scores were also predicted by their rsfMRI correlate scores. This structure was consistent across algorithms and independence test statistics.

## 4 Discussion

In this manuscript, we present a novel approach to study the interaction between behavioural variables and their neural correlates. We demonstrate this approach by investigating the relationship between autistic traits and their resting-state functional connectivity correlates identified through connectome-based predictive modelling^9^. Our results indicate that the network of associations between behavioural measures of autistic traits is not the same as the network of their neural correlates. We further demonstrate that complex causal interactions become apparent when behavioural traits and their neural correlates are considered together. This shows that the network-based regression approach can provide new insights into otherwise inaccessible aspects of brain-behaviour associations.

To determine if the results obtained through the CPM method with network-based regressors were reasonable, we compared the findings to the results obtained using traditional approach to studying brain-behaviour relationships. Regarding the connections implicated in each domain of the autistic traits, the current analysis indicated that lower connections strength of the frontoparietal control network (FCPN) was associated with higher difficulties across all autistic trait domains. This result is consistent with other reports that suggest a central role of the FCPN in autism^25,26^, possibly due to a shared influence of executive function difficulties mediated by the FCPN on all domains of autistic traits^27^. Further, a weaker connection between the default mode and the FCPN was associated with greater difficulties across domains. Differences in DMN connectivity is the most consistent findings in autism neuroimaging research^28^. Altered DMN connectivity may be linked to all autistic behaviours via a shared mechanism. For instance, this shared mechanism may lie in differences in generating and accessing internal representations that are mediated by the default mode network^29^. For instance, these internal representations may be necessary to adjust one’s behaviour, e.g. suppressing the urge to engage in repetitive behaviours, and to understand and interact with other people, e.g. by referring to previous experiences. Furthermore, the overlap in neural substrates implicated across autistic traits may arise from the simultaneous alterations in several large-scale networks as suggested by the triple network model^30^. According to this account, autistic traits may arise when differences in the DMN, salience network, and FCPN coincide. The current analysis also indicates that variation in the connectivity of at least two of these three networks (DMN, FCPN) is associated with all facets of autistic traits. In conclusion, the findings using the CPM method with network-based regressors are in line with previously implicated neural correlates of autistic traits.

Critically, our analysis using network-based regressors also identified brain correlates that were shared between only some of the behavioural traits. Communication and RRBIs were both associated with greater connectivity between the salience and the visual network. Greater connectivity of the salience network with sensory regions has been repeatedly reported in autism and may be related to sensory oversensitivity^31^ or increased reliance on visuospatial processing strategies^32,33^. The involvements of the same large-scale networks in these processes may explain their involvement in separate domains of autistic traits. Further, higher connectivity between the limbic network and the FCPN was associated with both higher difficulties with social interaction and RRBIs. Because the network-based regressors only contain the unique variance in each measure, any remaining overlap hints at a shared mechanism at the brain level. Thus, the brain-level association between social interaction and RRBI suggests that variation in both is driven by altered connectivity of the orbitofrontal cortex with the amygdala and striatum, possibly leading to both altered emotional processing and reward-related processing^34–36^.

Our results also indicated unique associations for each domain of autistic traits. Because the network-based regressors remove the overlap between the behavioural measures, these findings suggest unique brain-level mechanisms. For instance, higher connectivity of the somatomotor network with the FPCN and limbic network was only associated with higher communication difficulties but not the other autistic traits. Even in cases where the same large-scale network was implicated, the individual regions and connections differed between the autistic trait domains suggesting separation between the domains. This illustrates the utility of the network-based regression approach to identify unique mechanisms of behavioural traits.

The results obtained through network-based regression can be further employed to compare the network structure at the behavioural and brain level. This provides further indication about unique or shared mechanisms at each level of observation. In our autism example, there was a closer association between social and communication scores than between these scores and the RRBI score at the behavioural level. This result is consistent with the distinction between “social” and “non-social” traits suggested by some psychometric and genetic studies^37,38^ that is now incorporated in DSM-5. However, a different structure was indicated for the network of the rsfMRI correlates of the autistic traits. In the brain score network, correlates of social scores were associated with the correlates of RRBI scores, but not with the correlates of communication scores. This suggests that the close association of social and communication scores at the behavioural level is not due to a shared mechanism in brain function. In contrast, the association between the neural correlates of social and RRBI scores indicates that there is at least some overlap in the brain functional systems underlying these behaviours. In our analysis, Social and RRBI scores were both associated with differences in the connectivity of region within the FCPN, salience, and limbic network, while Communication scores were more associated with differences in the connectivity of the visual network. We further explored causal interaction within the networks. This analysis indicated complex causal interactions between behavioural measures of autistic traits and their neural correlates that could not be predicted from the network at either the behavioural or functional brain level alone. This causal network indicated that Communication scores mediate the association between RRBI and Social scores when accounting for the overlap in the associated functional brain systems. This shows that the network-based regression approach opens possibilities for analyses that contrast behavioural and brain-level associations to identify potentially shared mechanisms by using the information at the behavioural level to minimize any overlap in the regressors.

To construct a network of neural correlates of behavioural traits, we had to address several methodological challenges. Namely, in the first step of the analysis, we identified the functional connectivity correlates of the unique variance in autistic traits in a large, openly accessible database. The principal challenge here is the high dimensionality of the functional connectivity data. Even with a relatively coarse parcellation of the brain, the number of features by far exceeds the number of participants. For instance, a parcellation with 100 ROIs produces 4,950 features, assuming that the investigator excluded the diagonal and identical features in a symmetric adjacency matrix. Several methods have been developed to tackle this problem. These methods include complex machine learning pipelines, e.g. HPC Netmats Metatrawls, and dimensionality reduction approaches, such as partial least-squares analysis (PLS)^39^ or canonical correlation analysis (CCA)^40^. In the current analysis, we employed the connectome-based predictive modelling (CPM) method^9^ for its relative simplicity and clear interpretability^41^. This method identifies functional connectome features that are most closely associated with the behavioural variable of interest and mitigates overfitting through straightforward cross-validation. Despite its simplicity, the method has been successfully employed to identify functional connectome correlates in several studies^42–44^. Even though the modelling approach is relatively simple, the impact of several methodological choices still needs to be evaluated. Our results show that the choice of the functional connectivity metric, the resolution of the parcellation scheme, and the inclusion of global signal regression strongly impact on the strength of association between the functional connectivity features and the behavioural measures. We recommend that researchers who wish to apply the brain-behaviour network approach evaluate and report the impact of these methodological choices on the strength of the brain-behaviour association.

The results of the current analysis show that connectome-predictive modelling (CPM) can identify functional connections that are associated with each facet of the autistic trait triad. The summed brain scores significantly predicted the behavioural ratings and explained between 20 and 27% of the variance. The proportion of explained variance is lower than in studies that predicted cognitive task performance^43^ but is similar to the explained variance in a large-scale study with questionnaire measures (see https://db.humanconnectome.org/megatrawl/index.html). Measurement considerations aside, the explained variance in the current analysis may also be lower due to the variability in the dataset. We utilised data from ABIDE and ABIDE-II, which are retrospective collections that were acquired without prior harmonisation of the protocol^16,17^. Consequently, there is considerable variation between the acquisition sites^45^. We reduced this variability by applying stringent criteria for participant inclusion and by regressing the effect of extraneous variables from the connectome edges, but the remaining unaccounted variance is likely to have impacted on the amount of variance that could be explained in the behavioural measures.

It has to be noted that the network-based regression method has some limitations. First, relatively large samples are needed to obtain stable estimates of the networks. We used summed scores in our example to reduce the dimensionality of the behavioural measures. However, this involves a trade-off. Analyses with more detailed measures show that there is considerable variation within each domain at the behavioural level. For instance, RRBIs are dissociable into factors of repetitive behaviours and insistence on sameness^46^. Summing these dissociable domains may impact on the network structure at the behavioural level and influence subsequent analyses. Researchers will need to decide on the appropriate degree of granularity in the behavioural measures when employing the network-based regression approach for their research questions.

To conclude, we present a novel method to investigate the network structure of behavioural measures and their neural correlates. This method uses the unique variance in the behavioural variables to identify neural correlates. Subsequently, the behavioural and neural variables can be treated as nodes to compare the network structure at the level of behaviour and brain function. We show how this approach can be used to investigate if similarities at the behavioural level may be driven by similar mechanisms at the functional brain level. The approach presented here aligns closely with the shift towards complex system analysis in clinical psychology^1^ and expands the current toolkit to levels of analysis beyond behavioural traits.

## Author contributions statement

J.B. and D.B. developed the analysis plan, J.B. carried out the analyses and wrote the manuscript. All authors reviewed the manuscript.

## Additional information

### Open Science

The analysis code can be accessed on the Open Science Framework website: [OSF Link](https://osf.io/my8g6/?view_only=85c585b0f4ee4abb8e700afd2693b8c3). The ABIDE and ABIDE-II data is available to download from here: [NITRC link](http://fcon_1000.projects.nitrc.org/indi/abide/).

