## Supplementary Materials for "More than the sum of its parts: Merging network psychometrics and network neuroscience with application in autism"

### 1 Supplementary Introduction

Autism spectrum condition (ASC) is characterised by atypicalities in social interaction and communication, alongside restricted and repetitive behaviours and narrow interests (RRBI). Traditionally, diagnostic criteria require difficulties in at least two of these three domains for a diagnosis (DSM-IV)[1] . However, there is some debate regarding the separation between the domains. On one hand, it has been noted that atypicalities in all three domains frequently co-occur[1], possibly reflecting a shared genetic or neurocognitive mechanism. On the other hand, the 'fractionable triad' account suggests that unique mechanisms contribute to difficulties in each domain[2]. Factor analyses of scores from diagnostic interviews or behaviour ratings are inconclusive. Several studies indicate that a large proportion of variance loads onto a single factor in principal component analysis[3,4], which is interpreted as support for a single factor underlying the triad of autistic behaviours. Yet, other studies found support for a factor structure with two, three, or more factors[5–9]. A meta-analysis indicated that studies that identified more than one factor, typically divided the social aspects (social interaction, communication) from the putatively less social aspects (RRBIs)[10]. However, a more recent study that used a hierarchical cluster model indicated that communication and RRBIs segregated together with social interaction as a separate factor[11]. Investigations into the genetic underpinning also suggest some separation between the triad of autistic traits. For instance, findings from twin studies suggest a unique genetic contribution to each facet of the autism triad. Associations between phenotypic measures of autistic traits were modest across the full-range of scores and in the impaired range (most extreme 5% of scores) in a sample of over 3,000 twin pairs both at 8 years and 12 years of age[12–14]. The analysis of heritability indicated that the autistic traits showed a modest genetic overlap and substantial genetic specificity. Further support for unique influences on each domain of the autistic triad comes from studies that investigated the broader autism phenotype in undiagnosed family members of autistic individuals. These studies indicate that family members may show some aspect of autistic behaviours but not others[15–17], e.g. difficulties in social interaction without restricted or repetitive behaviours. This suggests that autistic traits can segregate. Similarly, a recent cluster analysis indicated that subgroups with relatively milder difficulties in one domain of autistic traits can also be distinguished in autistic groups[18]. In summary, the behavioural and genetic evidence suggests that the domains of atypicality in autism (social interaction, communication, RRBIs) are linked but can be fractionated. This provides an ideal test case for a network approach that aims to capture the relation between associated variables.

Neuroimaging research has associated the autistic trait triad with partially distinct networks of brain regions. Social cognition associated with the social interaction aspect of the autistic trait triad is thought to be supported by a distributed network of brain regions, collectively referred to as the "social brain"[19]. This network comprises of the amygdala, the dorsal and ventral medial prefrontal cortices (dmPFC, vmPFC), the anterior and posterior cingulate cortex (ACC, PCC), the posterior superior temporal sulcus (pSTS), the temporoparietal junction (TPJ), the inferior occipital gyrus (IOG), the fusiform face area (FFA), and the insula. These regions showed increased responses in tasks that tap social processing, including the decoding of facial expressions[20] and theory-of-mind[21]. These social brain regions show reduced activation during social processing tasks in autism[22,23]. Further, the connectivity between regions of the social regions is weaker in autism, yet the connectivity with regions outside of the social brain is stronger compared to neurotypicals[24–26]. Differences in social brain connectivity are also apparent at rest. Assaf et al. 2010 reported reduced connectivity of the canonical default mode (DMN) and salience network (SN) that contain social brain regions. Reduced connectivity of the DMN and SN was related to the severity of social symptoms[27]. Communication atypicalities in autism have been linked to differences in classic language areas, such as Broca's and Wernicke's area. Individuals with autism show reduced activation in the left inferior frontal cortex (Broca's area) and increased activation in the superior temporal gyrus (Wernicke's area) in lexical and semantic processing tasks[28–30]. Differences in connectivity within the language network is also apparent at rest. The left inferior frontal cortex (Broca's area) has been found to show reduced connectivity in children and adolescents with autism, while the superior temporal gyrus (Wernicke's area) shows reduced connectivity in autistic adults[31]. Differences related to communication difficulties in autism are also observed outside of the core language network. This includes increased activation of homologous areas in the right hemisphere during language processing[25,28] and activations in areas typically involved in visuospatial processing[29,32], i.e. the lateral occipital cortex and inferior parietal sulcus. Similarly, studies that investigated functional connectivity at rest indicated an association of higher communication difficulties with increased connectivity across the whole brain 38, reduced connectivity of the DMN and SN[27,33], and increased connectivity of the lateral occipital cortex[34]. Repetitive and stereotyped behaviours and narrow interests (RRBIs) have been associated with differences in systems involved in sensation, motor control, and reward-related processing. Specifically, higher RRBIs scores are linked to greater connectivity between the striatum with occipital and frontal areas[35], alongside lower connectivity of the striatum with cortical motor and sensory areas[33,36]. Further, higher RRBIs scores are associated with lower connectivity of frontoparietal regions[36,37], broadly consistent with the DMN. Furthermore, salience network hyperconnectivity has been found to predict RRBIs scores in autistic children[38].

As is apparent from this summary, there is a rich literature on the brain correlates of autistic traits. In some cases, the

[1]Please note that we focus on DSM-IV definition because most of the research and the data that was used for the analysis used DSM-IV criteria. DSM-5 uses two domains of difficulties, i.e. social and non-social.

associations overlap. For instance, the default mode and salience network are implicated across all domains of the autistic triad. Other associations appear to be unique to each domain, e.g. the involvement of the basal ganglia in RRBIs. The interplay between brain systems that are associated with autistic behaviours may be key. For instance, the involvement of the same brain system may explain the overlap of characteristics at the behavioural level. However, the interaction between autistic traits and their neural correlates has so far not been investigated. By applying a network approach, we aim to fill this gap here. To this end, we made use of large databases that collected resting-state fMRI and characterised autistics traits with the same assessment instrument. Using these data, we applied the connectome-based predictive modelling (CPM) method[39] to identify the rsfMRI correlates of autistic traits. To characterise the relations between behavioural and neuroimaging data, we applied network psychometric and causal inference methods. Based on the psychometric literature on autistic traits summarized above, we expected a closer association between social and communication difficulties than between these traits with RRBIs. We did not have a strong expectation regarding the association between the neural correlates but assumed a similar structure at the neural as in the behavioural data by default.

### 2 Supplementary Methods

#### 2.1 fMRI processing

The data were processed using the standard configuration of the Configurable Pipeline for the Analysis of Connectomes (C-PAC) v. 1.6.2[40]. C-PAC is an automated state-of-the-art pipeline for reproducible processing of large-scale data. The current analysis was run using the singularity image distributed via the C-PAC website to process the data on a high-performance computing cluster. The configuration file for the pipeline is provided in the Appendix so that the results can be exactly reproduced. The full details of the processing pipeline are available from the C-PAC website (https://fcp-indi.github.io/docs/user/preprocessing). The following provides an overview of the preprocessing steps. For anatomical preprocessing, a non-linear transform between images and a 2mm MNI brain-only template were calculated using ANTs v2.2.0. The images were then skull-stripped using AFNI's *3dSkullStrip* and subsequently segmented into WM, GM, and CSF using FSL's *FAST* tool[41]. The resulting WM mask was multiplied by a WM prior map that was transformed into individual space using the inverse of the linear transforms calculated through ANTs. A CSF mask was multiplied by a ventricle map derived from the Harvard-Oxford atlas distributed with FSL[42]. Skull-stripped images and grey matter tissue maps were transformed into MNI space at 2mm resolution.

For functional preprocessing, motion correction was performed using a two-stage approach in which the images were first co-registered to the mean of fMRI sequence and then a new mean was calculated and used as the target for a second registration (AFNI *3dvolreg*[43]). A 7-degree of freedom linear transform between the mean fMRI and the structural image was calculated using FSL's boundary-based registration[44]. Nuisance variable regression (NVR) was performed on the motion-corrected data using a 2nd-order polynomial, a 24-regressor model of motion[45], 5 nuisance signals identified via principal components analysis of signals obtained from white matter (*CompCor*[46]), and the mean CSF signal. WM and CSF signals were extracted using the previously described masks after transforming the fMRI data to match them in 2mm space using the inverse of the linear fMRI-sMRI transform. The NVR procedure was performed twice, with and without the inclusion of the global signal as a nuisance regressor. The results of the processing strategies were both entered into the predictive model in the later stages of the analysis (see Figure 1). The residuals of the NVR procedure were bandpass filtered ($0.001\text{Hz} < f < 0.1\text{Hz}$), written into MNI space at 2mm resolution and subsequently smoothed using a 6mm full-width half-maximum (FWHM) kernel.

Then, the time series for the parcellation described by Schaefer et al. 2018 were extracted with 200, 300, 400, and 1000 parcels[47]. We chose this parcellation because it provides a better account of fMRI activations than previous data-driven parcellations and is based on a large representative database (N=1,489). We evaluated to the optimal resolution for the purpose of the current analysis in the predictive model. To ensure that all included regions were sufficiently covered in all participants, we calculated the mean functional image, thresholded and binarized it at 70% intensity, and calculated the overlap between the atlas ROIs and the resulting image in all participants. ROIs that had less than 50% overlap with the binarised intensity image in any participant were excluded from the analysis. This resulted in the exclusion of 9 ROIs for the 200 ROI atlas, 16 for the 300 ROI atlas, 24 for the 400 ROI atlas, and 68 for the 1,000 ROI atlas. The excluded ROIs were located at the frontal and temporal pole.

#### 2.2 Functional connectome construction

We calculated the functional connectome matrices using three approaches, namely correlation, partial correlation, and tangent-space embedding. For the correlation approach, we calculated the Pearson correlation between each pairwise combination of ROIs in the parcellation. For the partial correlation approach, the Pearson correlation of each pairwise combination of ROIs was calculated after regressing the effect of other ROIs from both time series. For the tangent space embedding approach, we used the method described by Hutchison et al. 2010[48] that models each participant as a deviation from the group average connectome. For all approaches, the implementation of the method in Nilearn v. 0.6.2 was used[49]. To reduce the influence of extraneous variables, we applied a similar regression approach to the Human Connectome Project Mega-Trawl analysis

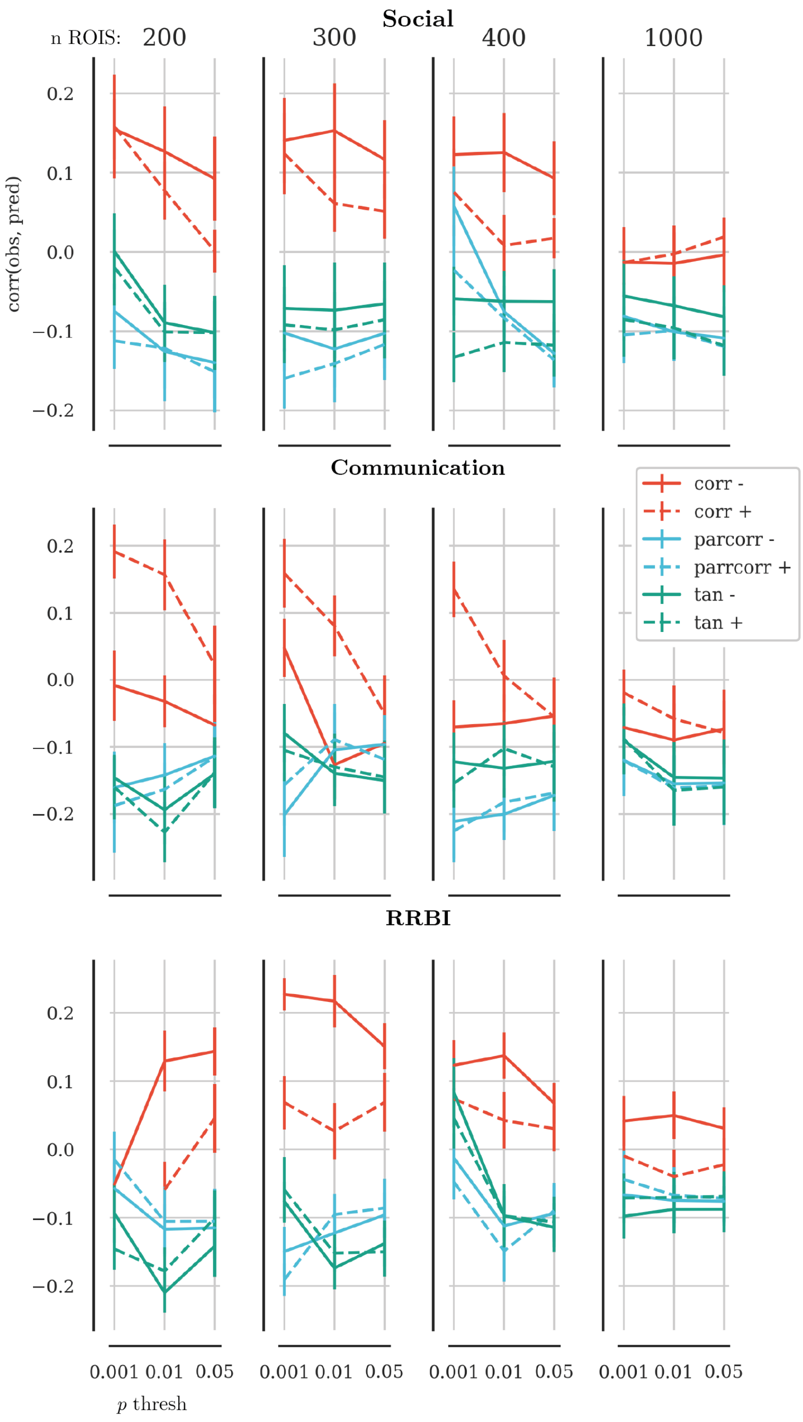


**Figure 1.** Results of the parameter tuning. The x-axis shows the *p*-value threshold used to select edges that were associated with behaviour ratings scores. The y-axis shows the correlation between the predicted scores and the observed scores in held-out data. Solid lines show results based on connectomes that were constructed without global signal regression, dashed lines indicate results based on connectomes with global signal regression. Red lines show results based on connectomes that used Pearson correlation as the edge definition, blue lines show the results for partial correlations, and green lines for tangent-space embedding. The panel in each row show the results for brain parcellations with 200, 300, 400, and 1,000 ROIs. The overall best results across behaviour scales was observed

(https://db.humanconnectome.org/megatrawl/HCP820_MegaTrawl_April2016.pdf). Namely, we regressed the effect of age, age2, acquisition site, framewise displacement, DVARS, intracranial volume (ICV), and total grey matter (GM) volume from each edge in the functional connectome in an ordinary least-squares regression model. ICV and total GM volume were estimated using FreeSurfer.

##### 2.3 Quality control

Functional MRI connectomics have been shown to be particularly sensitive to participant movement[50]. In order to mitigate the influence of motion, we applied a combination of steps. First, we only included participants with structural and functional data that were rated as useable by expert human assessors. These assessments are included in the phenotypic data file of the ABIDE and ABIDE-II database. For some sites, scans were assessed by multiple assessors. We only included a participant if all assessors rated the data as useable. Further, we evaluated the functional data using metrics calculated using the MRIQC pipeline[51]. We only included data that met conservative thresholds for framewise displacement (<0.5 mm[50]) and DVARS (<5%[52]). In the remaining participants, the effect of participant motion was mitigated by regression of nuisance signals, including noise components estimated in the white matter[46]. Furthermore, we regressed mean FD and standardised DVARS from each edge in the functional connectome. The quality metrics were not associated with social and communication scores (all $p>0.16$ uncorrected). RRBI scores were associated with DVARS ($p=0.032$ uncorrected), but not FD ($p=0.214$ uncorrected).
